# PRRSV evades innate immune cGAS-STING antiviral function via its Nsp5 to deter STING translocation and activation

**DOI:** 10.1101/2025.02.24.639812

**Authors:** Yulin Xu, Chenglin Chi, Qihang Xin, Jiang Yu, Yuyu Zhang, Pingping Zhang, Wangli Zheng, Sen Jiang, Wanglong Zheng, Nanhua Chen, Jiaqiang Wu, Jianzhong Zhu

## Abstract

Porcine Reproductive and Respiratory Syndrome Virus (PRRSV) is an important pathogen that seriously endangers pig breeding, causing significant economic losses to the global swine industry. Our previous study found that the DNA sensing innate cGAS-STING signaling pathway plays an important role in inducing interferon (IFN) upon PRRSV infection and inhibition of PRRSV replication. However, the immune evasion and its mechanism by PRRSV is still unclear. In current study, we found that PRRSV Non-structural protein 5 (Nsp5) strongly inhibits the cGAS-STING-IFN antiviral response. Further, we found that Nsp5 interacts with STING, blocking STING transport from ER to Golgi apparatus and interfering with STING recruitment of TBK1/IKKε/IRF3. Finally, we demonstrated that the Nsp5 36-47 and 58-67 amino acid regions are critical regions for inhibiting STING activity and PRRSV replication. The study describes a novel mechanism by PRRSV for suppression of the host innate antiviral response and have implications for our understanding of PRRSV pathogenesis.

**Author summary:** The innate immune system is the first line of host defense against infections. PRRSV is an immunosuppressive pathogen that has developed different strategies to evade the host innate immune responses. Here, we demonstrate for the first time that PRRSV utilizes its non-structural protein Nsp5 to inhibit the innate immune cGAS-STING-IFN signaling pathway, promoting the viral replication. Nsp5 can inhibit the cGAS-STING pathway by interacting with STING, blocking STING transport from ER to Golgi apparatus and interfering with STING recruitment of TBK1/IKKε/IRF3. In particular, the 36-47 and 58-67 amino acid regions of the Nsp5 is indispensable. This information will help us understand PRRSV-host interactions and provides new insights into the prevention and control of PRRSV.

## Introduction

Porcine reproductive and respiratory syndrome (PRRS) is an economically important viral disease all over the world characterized by reproductive failure in sows and respiratory diseases in growing pigs [1]. PRRS was first reported in the United States in 1987 and it was first identified in China in 1996 [2]. The pathogenic agent of PRRS is porcine reproductive and respiratory syndrome virus (PRRSV), a positive-sense single-stranded RNA virus. It belongs to the family *Arteriviridae*, and can be divided into two species, PRRSV-1 (the prototype European Lelystad virus LV strain) and PRRSV-2 (prototype North American VR2332 strain) [3]. PRRSV is an enveloped virus with a genome approximately 15.4 kb in length and contains at least 10 open reading frames (ORFs) encoding nonstructural polyproteins by ORF1a and ORF1b, and eight structural proteins (GP2, E, GP3, GP4, GP5, ORF5a, M, and N) by ORF2-7 [4]. The viral polyproteins are generated during infection by differential ribosome frameshift (RFS) events at two genomic sites, which include the pp1a, pp1a-nsp2N, pp1a-nsp2TF and pp1ab. These polyproteins are cleaved into at least 16 nonstructural proteins (Nsps) composed of Nsp1α, Nsp1β, Nsp2N, Nsp2TF, Nsp2 to Nsp6, Nsp7α, Nsp7β, Nsp8, Nsp9 to Nsp12, by viral proteases nsp1α, nsp1β, nsp2 and nsp4, collectively [4]. PRRSV nonstructural proteins play an important role in supporting viral replication and modulating host response [4–6].

The innate immunity serves as the first-line host defense against infections and utilizes multiple pattern recognition receptors (PRRs) to monitor the invading pathogens by sensing the pathogen-associated molecular patterns (PAMPs) and damage-associated molecular patterns (DAMPs) [7]. The innate immune PRRs include Toll-like receptors (TLRs), RIG-I-like receptors (RLRs), NOD-like receptors (NLRs), C-type lectin-like receptors (CLRs) and cytosolic DNA receptors (CDRs) [8–10]. Stimulator of interferon genes (STING) plays a pivotal role in innate immune responses as the central adaptor for DNA sensors, with cGAS as the most defined one [11, 12]. Upon DNA sensing, cGAS catalyzes the synthesis of second messenger 2’3’-cGAMP that activates STING in endoplasmic reticulum (ER) [12]. The activated STING transfers from ER to the trans-Golgi apparatus network where recruiting TBK1, IKKε, and subsequent transcription factor IRF3 to form a signalosome, by which both TBK1 and IKKε phosphorylate IRF3 [12–14]. The phosphorylated IRF3 then translocates into nucleus to activate the expression of antiviral type I interferons (IFNs), orchestrating innate immune responses [12].

STING has been shown to play a key role in sensing DNA virus infections, such as Herpes simplex virus 1 (HSV-1) [15], Human papillomavirus [16], Pseudorabies virus (PRV) [17], and African swine fever virus [18]. In addition, STING also plays an important role in the host’s resistance against RNA virus infections, such as Hanta virus [19], Zika virus [20] and PRRSV [21]. At the same time, viruses have evolved corresponding antagonizing machinery to evade the host’s antiviral immune defense. PRV utilizes its tegument protein UL13 to recruits RNF5 and inhibit STING-mediated antiviral immunity [17]. Zika virus evades STING antiviral response by cleaving cGAS via NS1-caspase-1 axis [20]. 3CL of SARS-CoV-2 inhibits K63-ubiquitin modification of STING to disrupt assembly of the STING functional complex and downstream antiviral signaling [22]. PRRSV as an RNA virus, has been shown to develop multiple mechanisms to antagonize RNA sensing innate immune antiviral response [5, 6]. However, it remained largely unclear whether and how PRRSV evades the DNA sensing cGAS-STING antiviral pathway.

In the present study, we screened the PRRSV proteins for interacting with porcine STING, and found that the viral Nsp5 was able to interact with STING and suppress STING activation by deterring its translocation from ER and recruitment of downstream TBK1/IKKε/IRF3. These findings have implications for PRRSV pathogenesis and reveal potential targets for the development of therapeutic strategies against PRRSV.

## Results

### The PRRSV Nsp3, Nsp5 and E proteins interact with porcine STING

As an RNA virus, the PRRSV genome is mainly recognized by innate immune RNA sensing receptors; nevertheless, our previous study demonstrated that the innate immune DNA receptor cGAS-STING signaling pathway plays an important role in sensing and defense of PRRSV infection. However, whether and how PRRSV evades the DNA sensing cGAS-STING antiviral pathway remains largely unclear. Here we focused on porcine STING and sought to examine the STING interacting PRRV proteins for inhibitory effect on cGAS-STING antiviral signaling. The Co-IP assay screening of all PRRSV proteins showed that PRRSV proteins Nsp3, Nsp5 and E interact with porcine STING (Fig 1A-C). The reverse Co-IP (Fig 1D) and co-localization assay (Fig 1E) further confirmed the interactions. The results clearly showed that PRRSV Nsp3, Nsp5 and E proteins interact with porcine STING, and potentially play a role in immune evasion of STING antiviral signaling.

**Fig 1.**
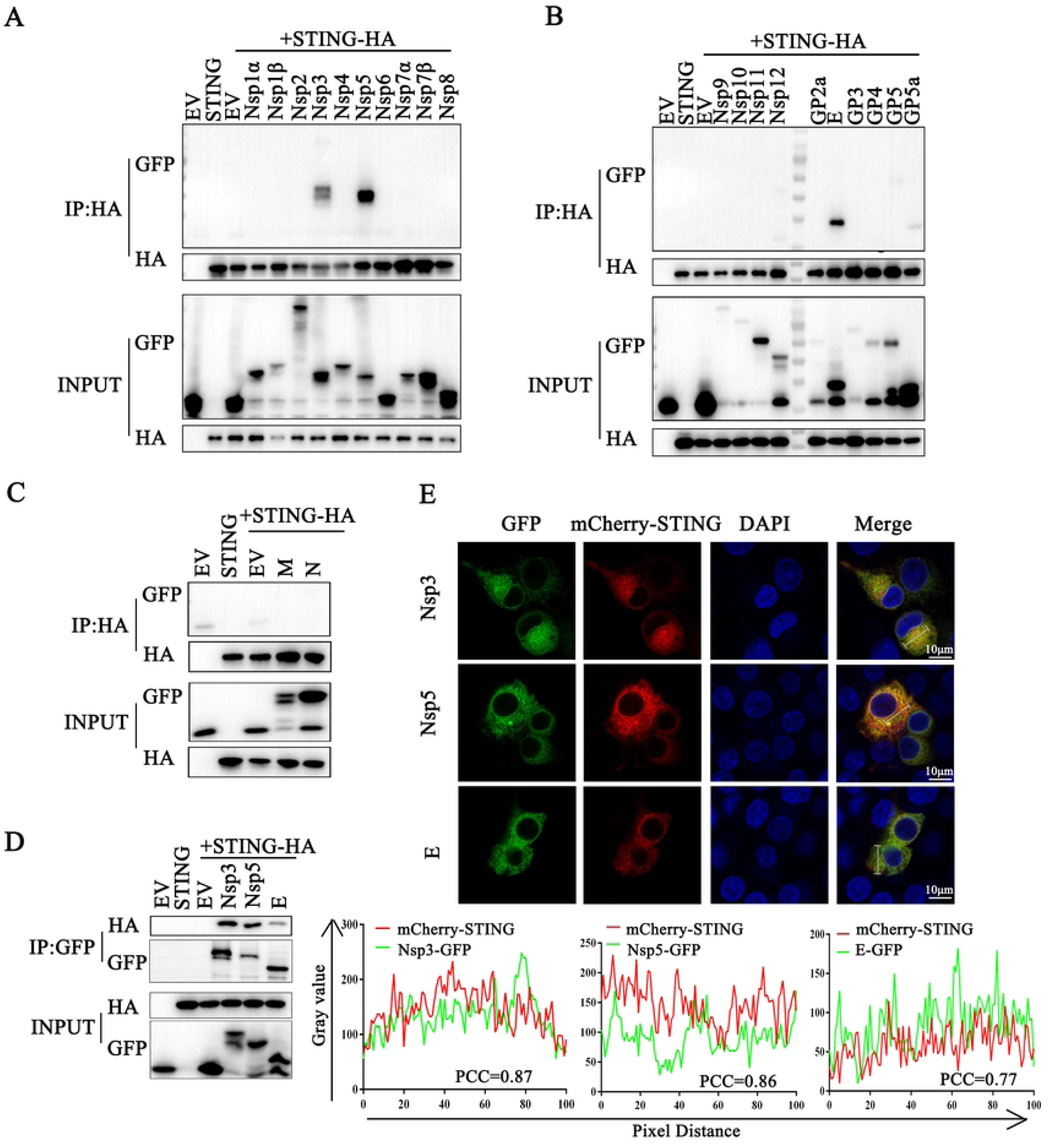
PRRSV proteins Nsp3, Nsp5 and E interact with STING. (**A-C**) HEK-293T cells were co-transfected pSTING-HA together with the indicated plasmids encoding GFP-tagged PRRSV viral proteins for 24 h. The cells were harvested for immunoprecipitation using rabbit anti-HA antibody, and IP samples and cell lysate inputs were subjected to Western blotting by using mouse anti-HA antibody and mouse anti-GFP antibody. (**D**) HEK-293T cells were co-transfected pSTING-HA together with the plasmids encoding GFP-tagged PRRSV viral proteins (Nsp3, Nsp5 and E) for 24 h, then harvested for immunoprecipitation using mouse anti-GFP antibody. The IP samples and cell lysate inputs were subjected to Western blotting by using rabbit anti-HA antibody and rabbit anti-GFP antibody. (**E**) 3D4/21 cells were co-transfected with mCherry-STING together with the indicated plasmid encoding GFP-tagged PRRSV viral proteins (Nsp3, Nsp5 and E) for 24 h, and examined for co-localization by confocal microscopy. The fluorescence intensity profile of mCherry (red) and GFP (green) was measured along the line, and the level of co-localization was quantified by calculating Pearson’s correlation coefficient (PCC) using ImageJ software. Scale bars of 10 µm are shown in the images.

### The PRRSV protein Nsp5 inhibits the cGAS-STING-mediated type I IFN response

Further, we evaluated innate immune modulation by PRRSV Nsp3, Nsp5 and E proteins in a luciferase reporter assay. We observed that Nsp3 cannot inhibit the promoter activity of the ISRE, IFN-β and ELAM mediated by the cGAS-STING pathway (Fig 2A-C). Nsp5 inhibits the activity of both ISRE and IFN-β promoters (Fig 2D-F), whereas E protein only inhibits the activity of ISRE promoter, but does not inhibit the activity of IFN-β and ELAM promoters (Fig 2G-I). Therefore, we selected the PRRSV Nsp5 for subsequent experiments.

**Fig 2.**
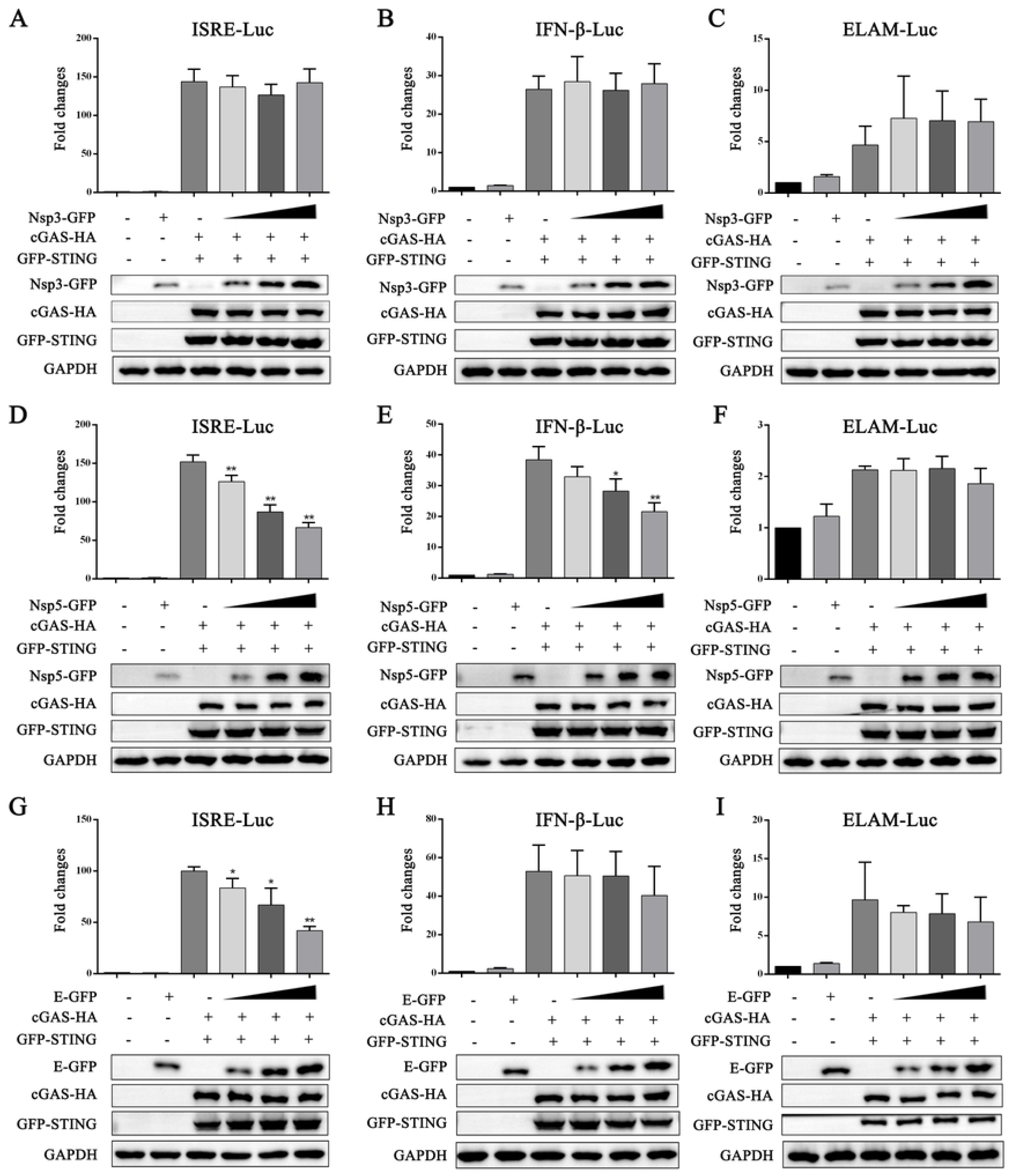
PRRSV Nsp5 can inhibits the activity of the ISRE and IFN-β promoters mediated by the cGAS-STING pathway. HEK-293T cells grown in 96-well plate (3×10^4^ cells/well) were transfected with ISRE (**A, D** and **G**), IFN-β (**B, E** and **H**), or ELAM (**C, F** and **I**) (10 ng) luciferase reporter, *Renilla* (0.2 ng) reporter, and plasmids expressing cGAS (10 ng) and STING (10 ng), together with increase amounts (20, 50, and 70 ng) of plasmids expressing Nsp3, Nsp5 and E. The total DNA was normalized to 100 ng/well and the luciferase activities were measured at 24 h post-transfection. The expressions of cGAS, STING, Nsp3, Nsp5, E and GAPDH were detected by Western blotting. * *p* < 0.05, ** *p* < 0.01.

In order to explore the impact of Nsp5 on the DNA sensing cGAS-STING pathway, we analyzed the effects of Nsp5 on the activation of relevant signaling proteins in the cGAS-STING signaling pathway using Western blotting. In transfected HEK-293T cells, the exogenous porcine cGAS-STING activated phosphorylations of TBK1 (p-TBK1) and IRF3 (p-IRF3) were both inhibited by Nsp5 in a dose dependent manner (Fig 3A). In porcine macrophage 3D4/21 cells, the levels of p-TBK1 and p-IRF3 induced by endogenous STING activation were also inhibited by Nsp5 in a dose dependent way (Fig 3B). Further, Nsp5 inhibited the IRF3 nuclear translocation upon STING activation by 2’3’-cGAMP (Fig 3C). These results demonstrated that PRRSV Nsp5 invades STING-mediated type I IFN signaling.

**Fig 3.**
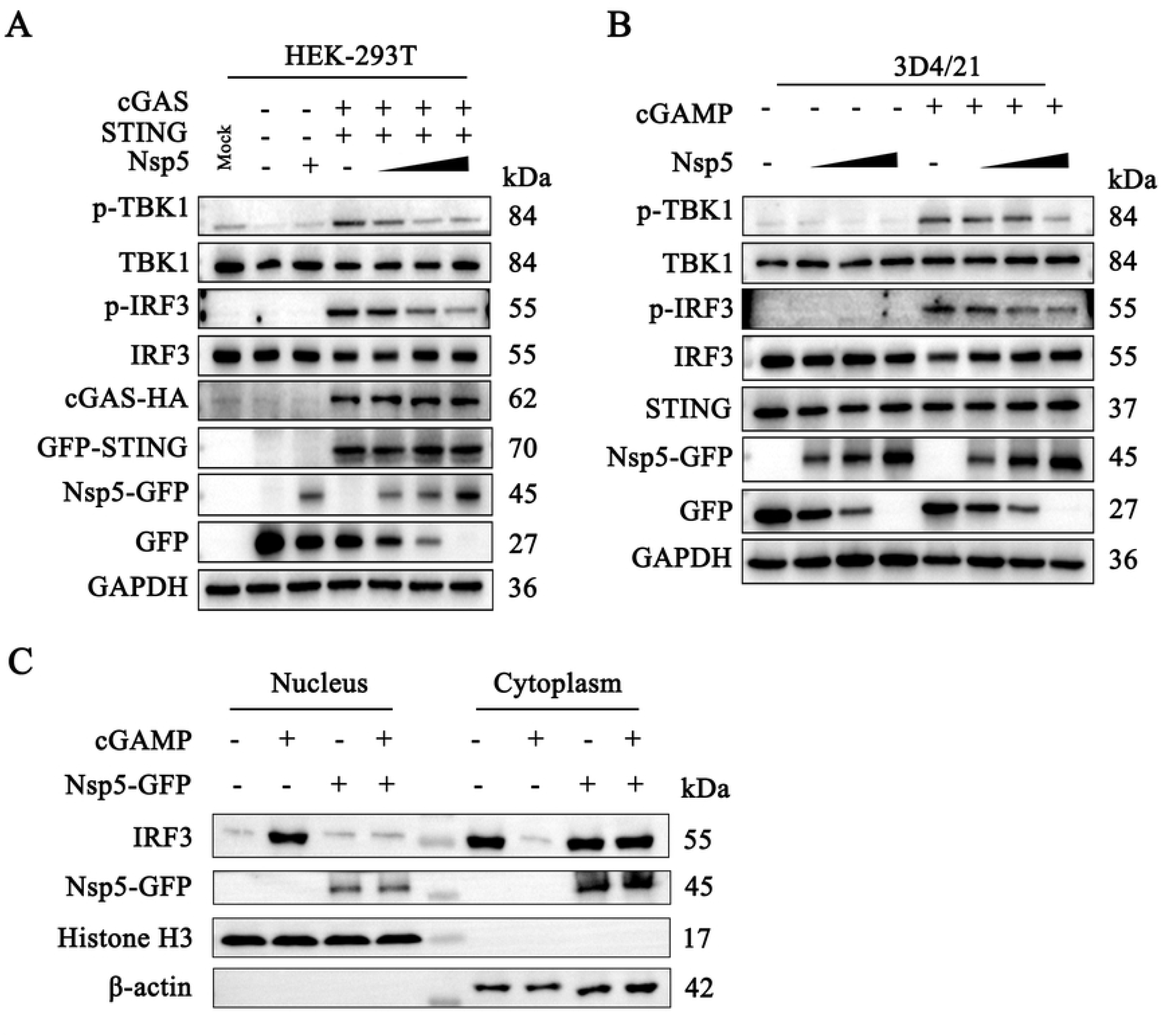
PRRSV Nsp5 inhibits the STING-mediated downstream signaling pathway. (**A**) HEK-293T cells in 6-well plate were transfected with cGAS (0.5 μg), STING (0.5 μg) and Nsp5 (0.25, 0.5 or 1 μg) for 24 h. Empty vector was added to keep the total DNA amount constant. (**B**) 3D4/21 cells in 12-well plate were transfected with Nsp5 (0.25, 0.5 or 1 μg) for 12 h, then cells were transfected with 2’3’-cGAMP (200 ng/mL) for 12 h. The cell lysates were subjected for Western blotting using the indicated antibodies. (**C**) 3D4/21 cells were transfected or not with Nsp5 (1 μg) for 24 h and then treated with or without 2’3’-cGAMP (1μg/mL) for 12 h. Cells were then harvested for extraction of cytoplasmic and nuclear proteins. The extracted proteins were determined by Western blotting with antibodies against IRF3, GFP, histone H3, and β-actin, respectively.

### PRRSV Nsp5 inhibits STING signaling by retaining STING at the ER

Our recent study showed that STING recruits both TBK1 and IKKε during translocation from ER to Golgi apparatus, and the STING, TBK1, IKKε and IRF3 may form a signalosome by which both TBK1 and IKKε phosphorylate IRF3 and other substrates for type I IFN response [13]. First, the interactions between porcine STING and three signaling proteins porcine TBK1, IKKε and IRF3 were examined by Co-IP assay. As expected, STING was able to interact with TBK1, IKKε and IRF3, respectively (Fig 4A-C). Although Nsp5 had interaction with STING (Fig 4D-E), but not with TBK1, IKKε and IRF3 (Fig 4F-H). However, Nsp5 was able to inhibit the interactions between STING and TBK1 (Fig 4I), between STING and IKKε (Fig 4J), between STING and IRF3 (Fig 4K), in dose dependent ways. More importantly, Nsp5 was able to inhibit the interactions between endogenous STING and TBK1/IKK**ε**/IRF3 in a dose dependent way (Fig 4L-N).

**Fig 4.**
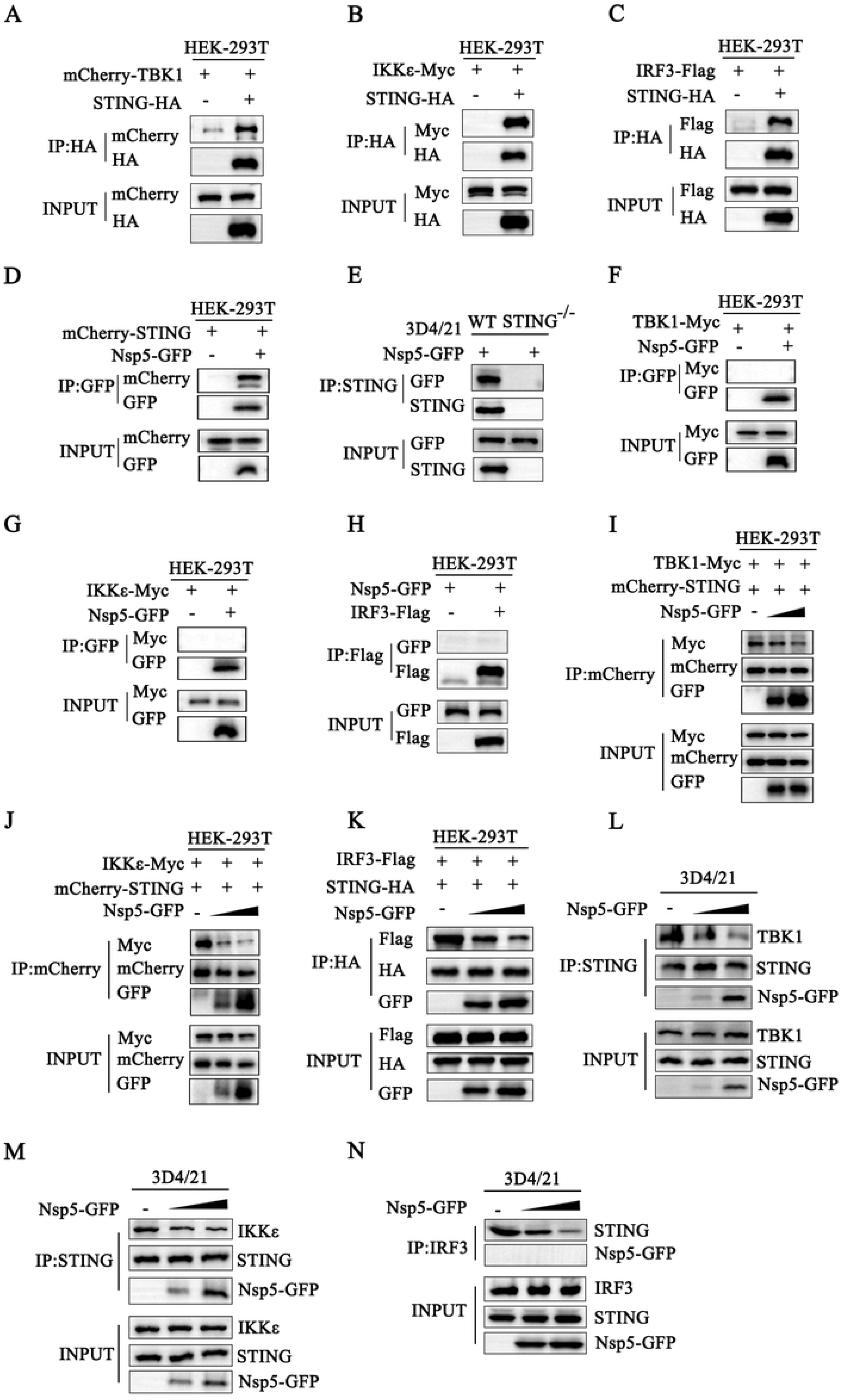
PRRSV Nsp5 inhibits the cGAS-STING pathway through its interaction with STING. (**A-C**) HEK-293T cells were cotransfected with mCherry-TBK1 and STING-HA, with IKKε-Myc and STING-HA, and with IRF3-Flag and STING-HA for 24 h. (**D**) HEK-293T cells were cotransfected with mCherry-STING and Nsp5-GFP for 24 h. (**E**) STING-deficient 3D4/21 cells were transfected with Nsp5-GFP for 24 h. (**F-H**) HEK-293T cells were cotransfected with TBK1-Myc and Nsp5-GFP (F), with IKKε-Myc and Nsp5-GFP (G), and with IRF3-Flag and Nsp5-GFP (H) for 24 h. (**I-K**) HEK-293T cells were cotransfected with 0.5 μg mCherry-STING, 0.5 μg TBK1-Myc, and increasing amounts of Nsp5-GFP (0.5 and 1 μg) for 24 h (I), with 0.5 μg mCherry-STING, 0.5 μg IKKε-Myc and increasing amounts of Nsp5-GFP (0.5 and 1 μg) for 24 h (J), with 0.5 μg STING-HA, 0.5 μg IRF3-Flag and increasing amounts of Nsp5-GFP (0.5 and 1 μg) for 24 h (K). (**L-N**) 3D4/21 cells were transfected with Nsp5-GFP (1 and 2 μg), then stimulated with 2’3’-cGAMP for 12 h. The cells were harvested and subjected for Co-IP and Western blotting using the indicated antibodies.

STING is an ER-resident protein, and its translocation from the ER to the Golgi apparatus is a hall marker of STING activation [23]. Considering that Nsp5 inhibits the cGAS-STING pathway through its interaction with STING and its interference of the recruitment of TBK1, IKKε and IRF3, we hypothesized that Nsp5 may interfere with STING translocation from ER to Golgi apparatus. Therefore, the ectopic Nsp5-GFP and mCherry-STING were expressed in porcine macrophages 3D4/21, followed by stimulation with 2’3’-cGAMP, and then cells were processed for immunostaining to detect the STING location. As predicted, STING was located at the ER of unstimulated cells, and upon stimulation, it transported from the ER to Golgi apparatus (Fig 5A), forming large puncta in Golgi apparatus (Fig 5B). In contrast, in Nsp5 expressed cells, STING remained in the ER and had no puncta formation upon 2’3’-cGAMP stimulation (Fig 5A-B). These data indicated that Nsp5 inhibits STING signaling by hindering STING trafficking.

**Fig 5.**
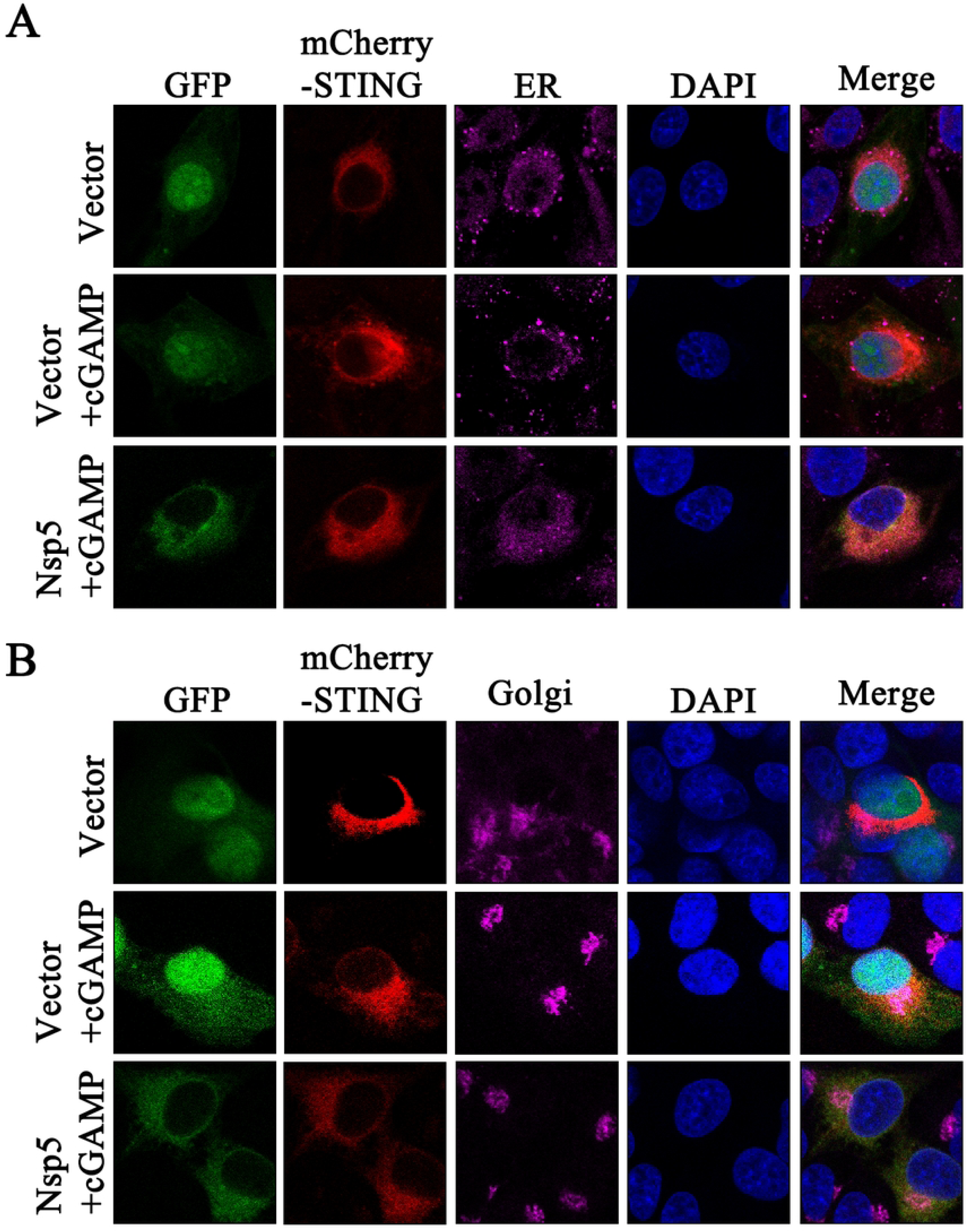
PRRSV Nsp5 blocks STING trafficking from the ER to the Golgi apparatus. (**A** and **B**) 3D4/21 cells were cotransfected with Nsp5 and STING expression plasmids for 24 h, and then treated with or without 2’3’-cGAMP for 12 h. Cells were fixed, permeabilized, and incubated with antibodies specific for Calreticulin (ER marker, A) and GORASP2 (Golgi marker, B), followed by counter-staining with DAPI. Samples were examined under a confocal microscope.

### Porcine STING 1-153 amino acids is a critical region for interaction with Nsp5

To identify which domain of porcine STING is important for STING interaction with Nsp5, five truncated mutants of STING were prepared (Fig 6A). Subsequently, Co-IP assay was used to identify the STING domains important for Nsp5 interaction. Domain mapping revealed that Nsp5 interacts with STING 1-339 and STING 1-190 (Fig 6B-C), but not with STING 191-378 and STING 153-339 (Fig 6D). Based on these results, we inferred that the STING 1-153 is the key domain for STING interaction with Nsp5. As proposed, the STING 1-153 is necessary and sufficient for STING interaction with Nsp5 (Fig 6E). Corresponding with these Co-IP results, the mCherry-STING 1-153 was co-localized very well with Nsp5-GFP in the cytoplasm of transfected cells (Fig 6F).

**Fig 6.**
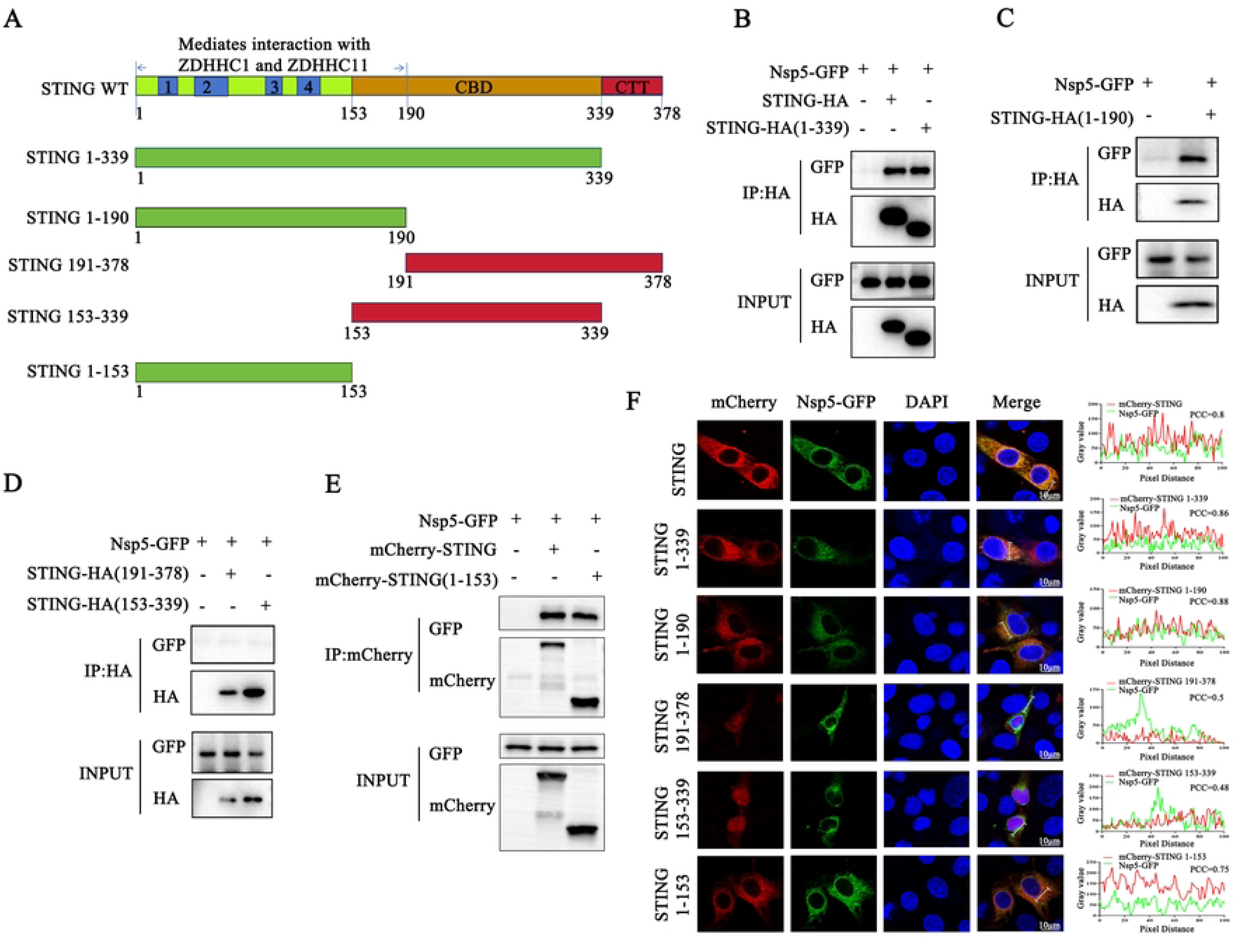
Porcine STING 1-153 amino acids is a critical region for interaction with Nsp5. (**A**) schematic of porcine STING molecular structure and its truncated mutants. (**B-E**) HEK-293T cells were cotransfected with STING or its truncation mutants and Nsp5-GFP for 24 h, and the cells were harvested and subjected for Co-IP and subsequent Western blot analysis using indicated antibodies. (**F**) mCherry-STING or its truncation mutants were cotransfected with Nsp5-GFP into 3D4/21 cells for 24 h. Cells were fixed, permeabilized, and examined for co-localization by confocal microscopy. The levels of co-localization between STING or its truncation mutants and Nsp5 were quantified by PCC using ImageJ software. Scale bars are10 µm.

### The 36-47 and 58-67 amino acids of Nsp5 are the key regions for interaction with STING

To identify the region where Nsp5 interacts with STING, we prepared ten truncated Nsp5 mutants: 1-101, 1-67, 1-57, 1-47, 1-35, 102-170, 68-170, 58-170, 48-170 and 36-170 (Fig 7A). The Co-IP assay results revealed that STING interacts with Nsp5 1-101 and 1-67, but not with Nsp5 68-170 and 102-170 (Fig 7B). Further, STING did not interact with Nsp5 1-57, 1-47 and 1-35 (Fig 7C). These data indicated that the 58-67 amino acids of Nsp5 is a necessary region for the interaction with STING. On the other hand, STING interacted with Nsp5 36-170, but not with Nsp5 48-170 and 58-170 (Fig 7D). The results indicated that 36-47 amino acids of Nsp5 is also a necessary region for the interaction with STING.

**Fig 7.**
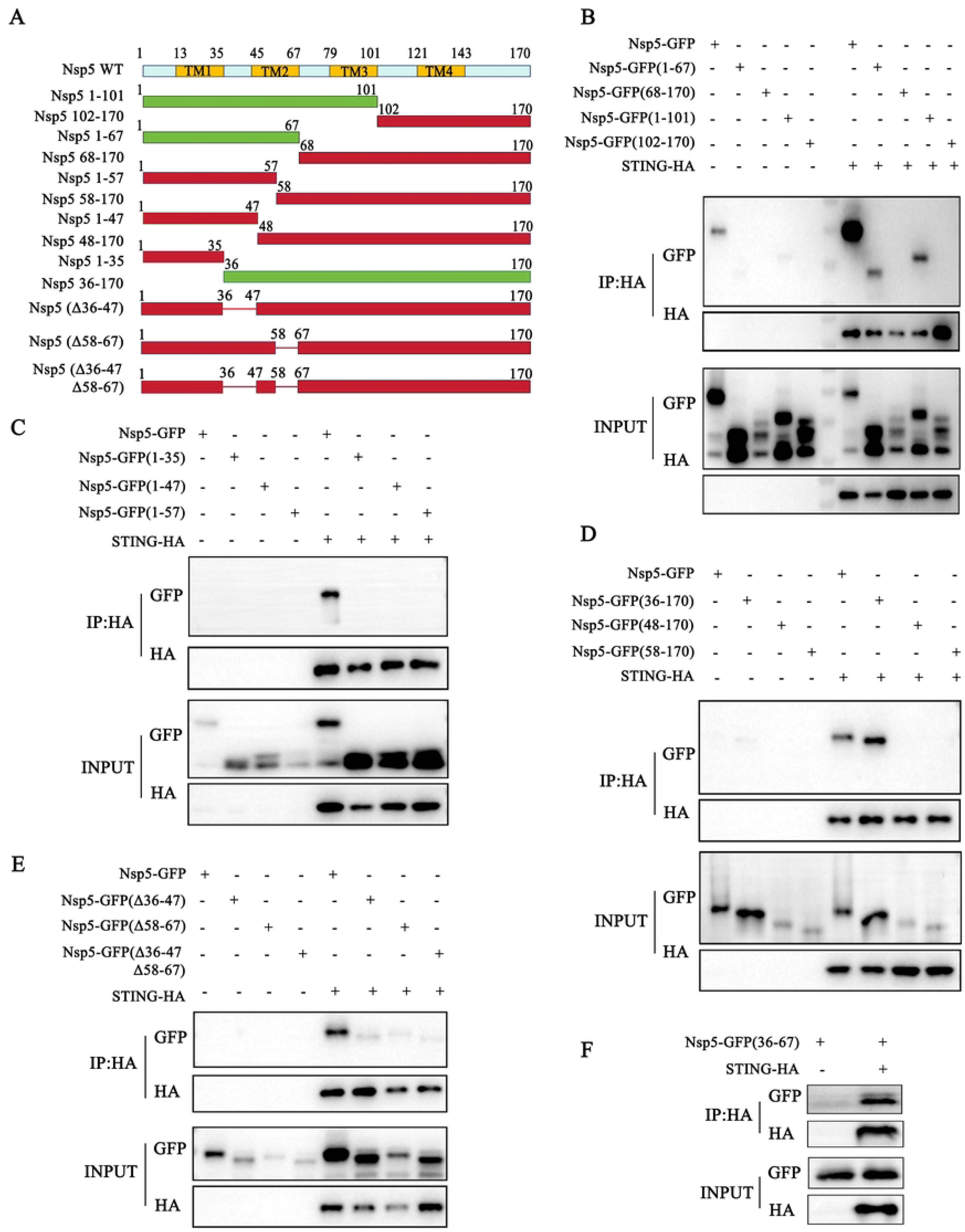
The 36-47 and 58-67 amino acids of Nsp5 are the key regions for interaction with STING. (**A**) Schematic of PRRSV Nsp5 molecular structure and its truncation and delection mutants. (**B-F**) Nsp5-GFP or its truncation mutants/deletion mutants were cotransfected with STING-HA into HEK-293T for 24 h, and the cells were harvested and subjected for Co-IP and subsequent Western blot analysis using indicated antibodies.

To ascertain whether Nsp5 36-47 and 58-67 amino acid regions of Nsp5 are important for Nsp5 interaction with STING, two single and one double deletion mutants of GFP-tagged Nsp5 were prepared (Δ36-47, Δ58-67 and Δ36-47Δ58-67). Co-IP analysis showed that three Nsp5 deletion mutants lost the interaction with STING (Fig 7E), confirming that the 36-47 and 58-67 amino acid regions of Nsp5 are indeed necessary for interaction with STING. In addition, we constructed the 36-67 amino acid region of Nsp5 covering these two necessary regions, and the Co-IP results showed that STING interacts with 36-67 region of Nsp5 (Fig 7F), suggesting that the two regions are key regions for Nsp5 interaction with STING. Consistently, the confocal microscopy illustrated that the regions of 36-47 and 58-67 amino acids are important for Nsp5 co-localization and binding to STING (Fig S1).

### The 36-47 and 58-67 amino acid regions of Nsp5 are crucial for interfering with STING signaling and PRRSV replication

We further investigated the impact of three deletions (Δ36-47, Δ58-67 and Δ36-47Δ58-67) in Nsp5 on the recruitment of TBK1 and IKKε by STING. The Co-IP results showed that, compared with full-length Nsp5, three deletion mutants lost the ability to disturb the interaction of exogenous STING and TBK1 (Fig 8A), the interaction of exogenous STING and IKKε (Fig 8B). Consistently, three deletion mutants of Nsp5 also lost the ability to disturb the interactions of endogenous STING and TBK1/IKKε (Fig 8C-D). Subsequently, we determined the effect of three Nsp5 deletions on STING trafficking. Upon 2’3’-cGAMP stimulation, neither Nsp5 deletion mutant could block STING translocation from the ER to the Golgi apparatus (Fig 9A-B). Further, compared with the full-length Nsp5, all three deletion mutants lost the ability to inhibit the ISRE and IFN-β promoter activity induced by the cGAS-STING pathway (Fig 9C-D).

**Fig 8.**
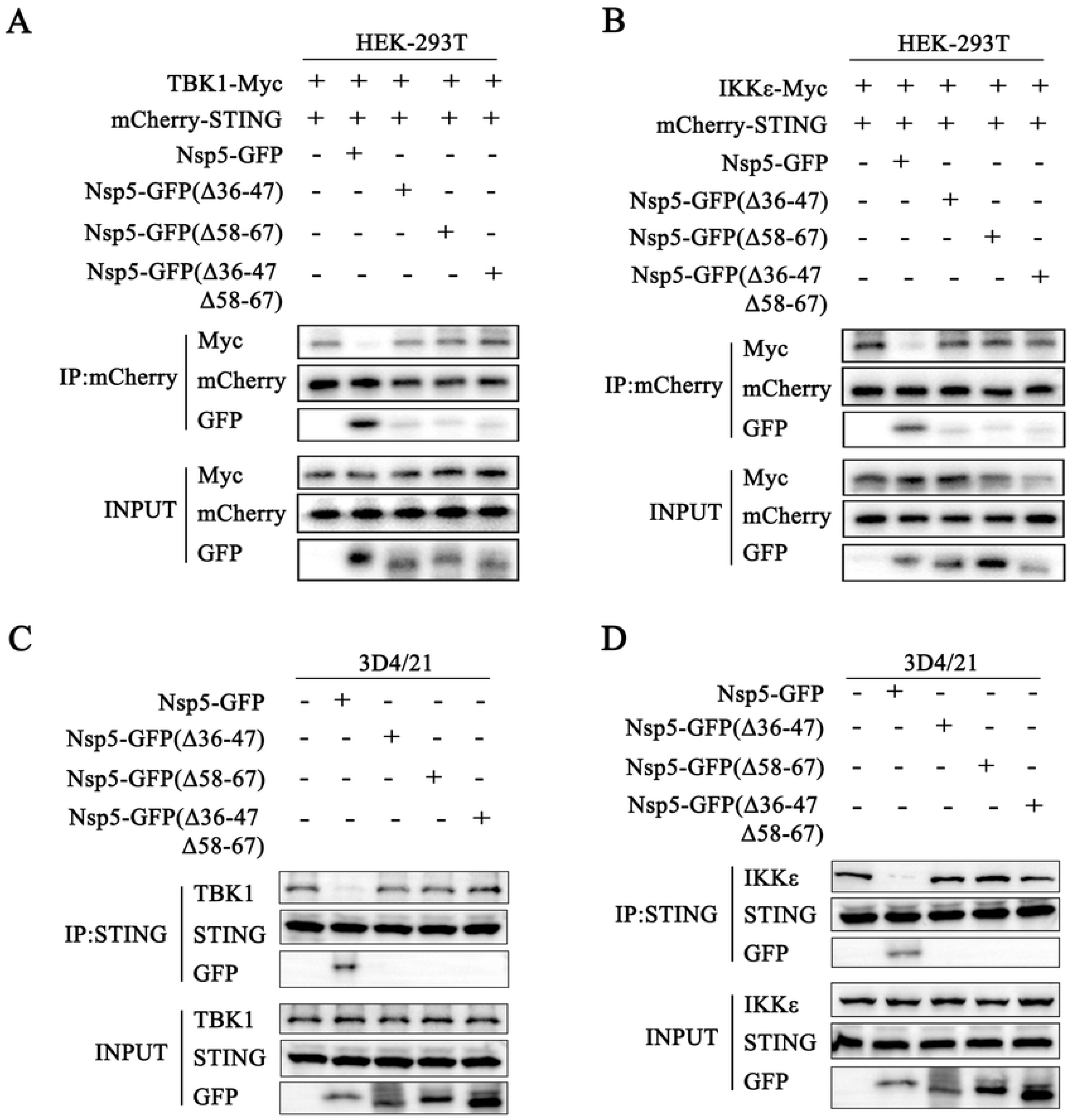
The regions of 36-47 and 58-67 amino acids of Nsp5 are crucial for interfering with STING signaling. (**A**) HEK-293T cells were cotransfected with TBK1-Myc together with mCherry-STING, Nsp5 or its deletion mutants for 24 h. (**B**) HEK-293T cells were cotransfected with IKKε-Myc together with mCherry-STING, Nsp5 or its deletion mutants for 24 h. Cell lysates were immunoprecipitated with anti-mCherry antibody and subjected for Co-IP and subsequent Western blot analysis. (**C-D**) 3D4/21 cells were transfected with Nsp5 or its deletion mutants for 24 h, and then stimulated with 2’3’-cGAMP for 12 h. Cell lysates were immunoprecipitated with anti-STING antibody and then analyzed by Western blotting for STING, TBK1, IKKε and GFP.

**Fig 9.**
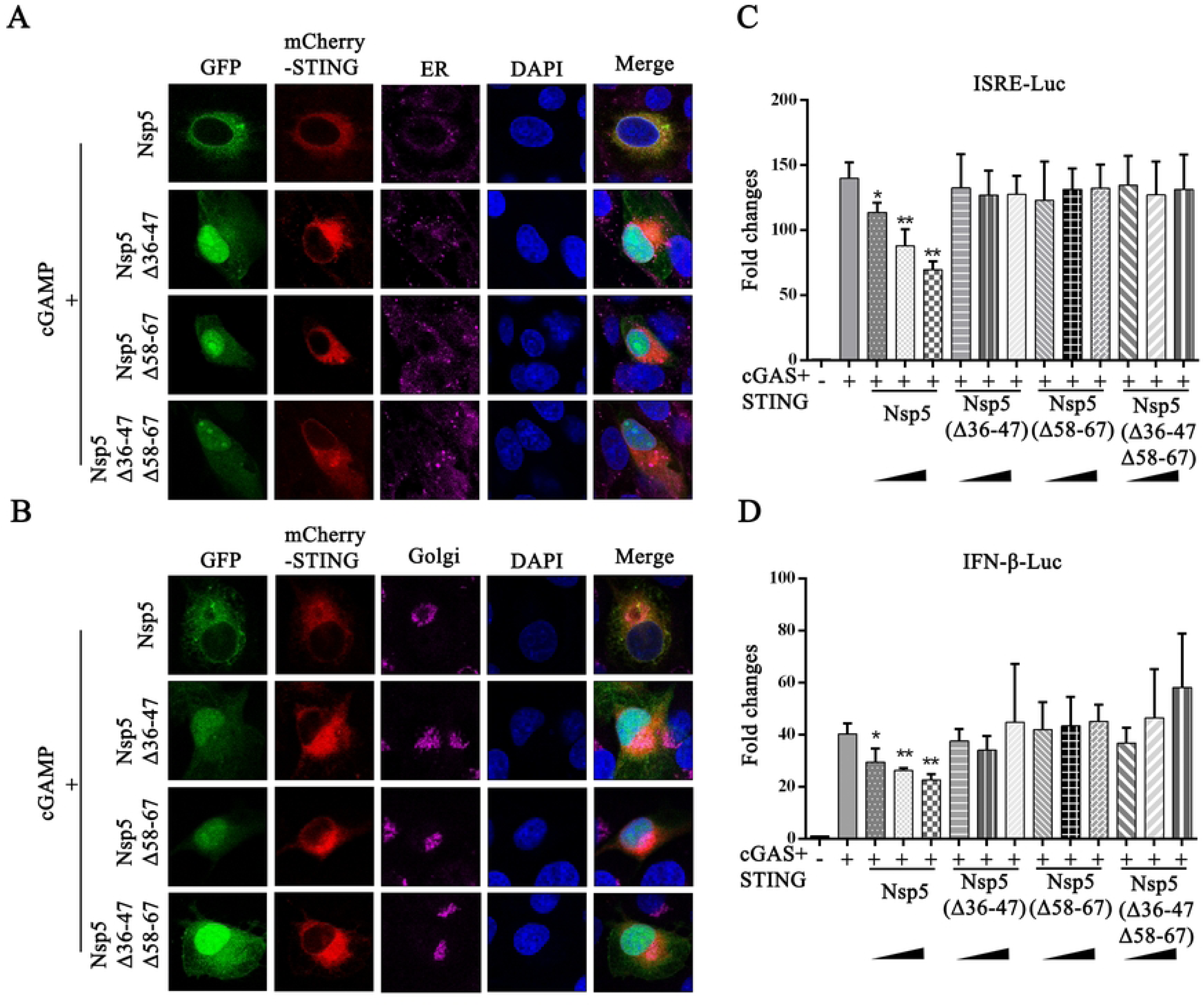
The regions of 36-47 and 58-67 amino acids of Nsp5 are crucial for blocking the STING trafficking and signaling. (**A-B**) 3D4/21 cells were cotransfected with STING and Nsp5 or Nsp5 deletion mutants and then treated with 2’3’-cGAMP. Cells were fixed, permeabilized and incubated with antibodies specific for Calreticulin (A) and GORASP2 (B), followed by DAPI staining. Samples were examined under a confocal microscope. (**C-D**) HEK-293T cells were transfected with the ISRE-Luc (C), IFN-β-Luc (D) reporter plasmid, together with Rluc reporter, cGAS, STING and Nsp5 or Nsp5 deletion mutant plasmids. Cells were collected 24 h after transfection, and ISRE/IFN-β promoter activity was evaluated via a luciferase reporter assay. * *p* < 0.05, ** *p* < 0.01

To further clarify the importance of the 36-47 and 58-67 amino acids of Nsp5 in PRRSV infection, we also tried to make the recombinant PRRSVs carrying the three deletion mutations by mutagenesis of infectious cDNA clones (Fig S2A-B). Cytopathic effects were observed at 48 hours post infection (hpi) in the recombinant PRRSV XJ17-5 (rXJ17-5) and recombinant PRRSV XJ17-5-EGFP (rXJ17-5-EGFP) infected Marc-145 cells but not in cells mock-infected and infected with recombinant PRRSVs carrying any deletion mutations (Fig S2C). In addition, the GFP signal was visualized under fluorescence microscope at 48 hpi in cells infected with the rPRRSV XJ17-5-EGFP but not in cells mock-infected and infected with rPRRSV XJ17-5 and recombinant PRRSVs carrying deletion mutations (Fig S2C). Together, these results demonstrated the critical roles of 36-47 and 58-67 amino acids of Nsp5 not only in blocking STING antiviral signaling, but also in PRRSV infection.

## DISCUSSION

As the RNA virus, PRRSV was thought to be recognized by innate immune RNA sensing receptors TLR3/7/8/9 and RLRs [24]. In our previous study, we systemically dissected the anti-PRRSV activities of nine porcine innate immune signaling adaptors [25]. The results indicated that multiple porcine PRR signaling pathways might be involved in the sensing of and fighting against PRRSV [25]. Further, we and another group clearly demonstrated that DNA sensing cGAS-STING pathway plays an important role in sensing and controlling PRRSV infection [21, 26]. However, the interplay between PRRSV and DNA sensor-related host defense is intricate and the viral evasion mechanism remains incompletely understood. In the present study, we identified a previously unrecognized mechanism by which PRRSV inhibits porcine STING antiviral signaling via blocking its trafficking. This viral immune evasion is dependent on the viral protein Nsp5, which interacts with STING in ER to impair STING’s transport to the Golgi apparatus, and in turn affects downstream antiviral immunity. This work provides a better understanding of the immune escape and pathogenesis mechanism of PRRSV.

During PRRSV infection, there is a complex relationship between host immune defense and viral infection [27–29]. Our results suggested that PRRSV Nsp5 can inhibit the phosphorylations of both TBK1 and IRF3 activated by cGAS-STING signaling. Nsp5 inhibits the activation of the cGAS-STING-IFN pathway by interacting with STING in ER, blocking STING transport from ER to Golgi apparatus and interfering with STING recruitment of TBK1/IKKε/IRF3. Interestingly, a recent research report showed that the Nsp2 of PRRSV impedes STING translocation from the ER to the Golgi apparatus by deubiquitinating STIM1 via Nsp2 deubiquitinating (DUB) activity [30], thereby facilitating immune escape. We did not see the interaction between Nsp2 and STING in our initial screening experiments, which may be due to various experimental conditions. Nevertheless, it is possible that several PRRSV non-structural proteins (Nsps) are involved in blocking STING trafficking and inhibiting STING antiviral signaling. Further exploration is needed to give deep insights into evasion of STING traffic and signaling by PRRSV.

There are relatively few studies on Nsp5, which is a conserved non-structural protein of PRRSV composed of approximately 170 amino acid residues [31]. Nsp5 is a hydrophobic transmembrane protein and can form a membranous structure in the cytoplasm that could be the site for viral replication [1]. Nsp5 participates in the formation and maintenance of double-membrane vesicles (DMV) during PRRSV infection, thereby promoting the synthesis and replication of viral RNA [32]. Also, Nsp5 promotes the formation and accumulation of cell autophagosomes [33, 34]. In terms of immune regulation, Nsp5 antagonizes JAK/STAT3 signaling by inducing degradation of STAT3 [35], and it also inhibits RLR signaling pathway by degrading multiple proteins of this pathway [36]. Therefore, Nsp5 can not only antagonize RNA sensor mediated innate immune response, but also DNA sensor mediated innate immune response.

In this study, we identified two key regions of Nsp5 for interacting with STING and attempted to make recombinant viruses carrying the Nsp5 deletion mutations by mutagenic infectious cDNA clones. However, it failed to rescue the recombinant viruses, likely due to the large number of deleted amino acids, highlighting the critical role of Nsp5 for PRRSV replication. Currently, we are attempting to identify key amino acids for interacting with STING and rescue recombinant viruses carrying point mutations in Nsp5, which will be useful to investigate the impact of Nsp5 on innate immunity in *vitro* and *vivo*.

In conclusion, this study revealed that PRRSV Nsp5 can inhibit DNA innate immune cGAS-STING antiviral signaling pathway, by blocking porcine STING trafficking from ER to Golgi apparatus and interfering STING recruitment of TBK1/IKKε/IRF3 (Fig 10). Importantly, the 36-47 and 58-67 amino acids of Nsp5 are the key regions for interacting with porcine STING, immune escape and PRRSV replication. These findings provide new insights into the pathogenesis mechanisms PRRSV and development of novel antiviral strategies against PRRSV.

**Fig 10.**
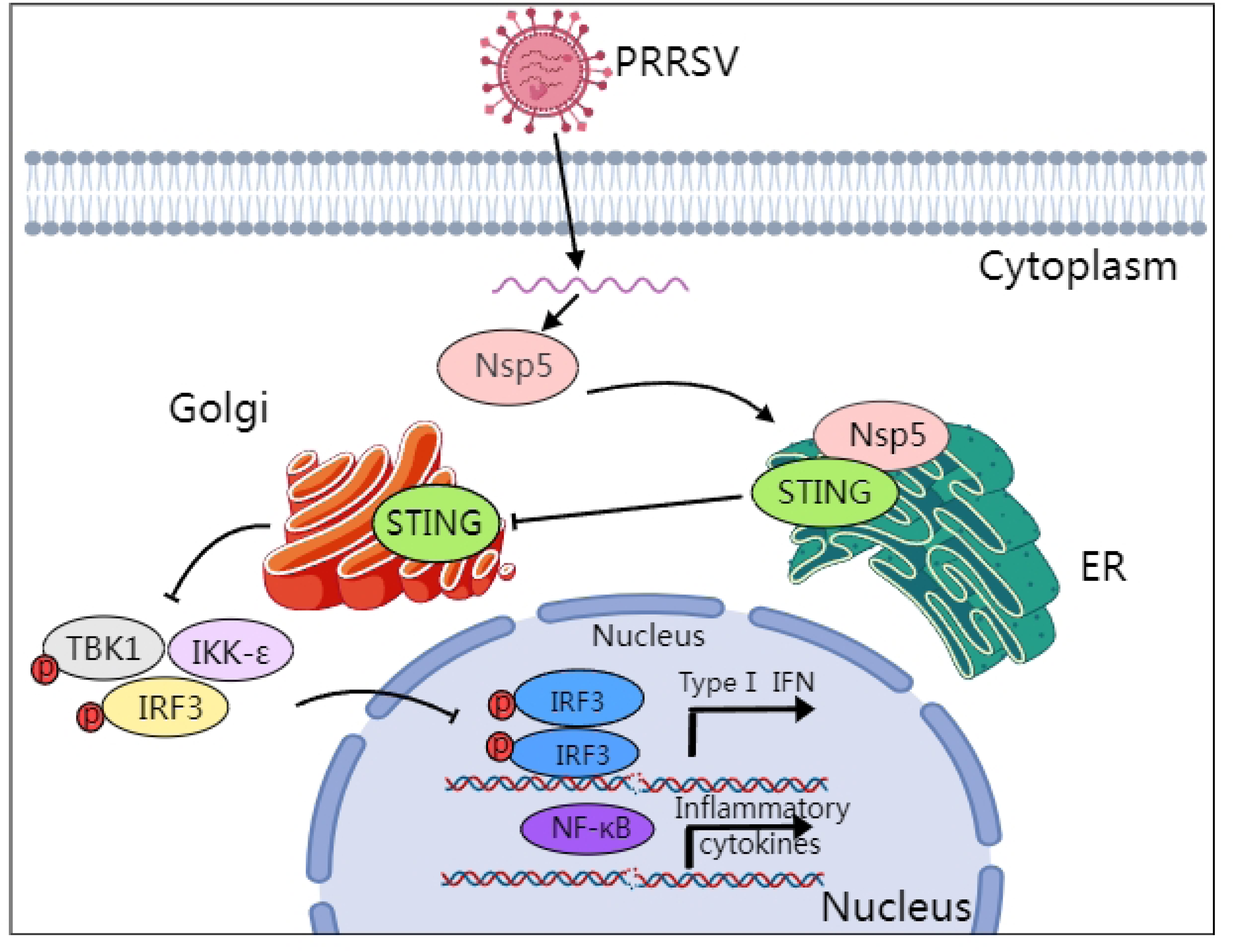
Schematic model of the mechanism by which PRRSV Nsp5 inhibits the cGAS-STING signaling pathway, created in MedPeer. The interaction between PRRSV Nsp5 and STING hinders the transport of STING from ER to Golgi apparatus, and thus disrupts the assembly of the STING/TBK1/IKKε/IRF3 signaling complex and downstream antiviral IFN response.

## Materials and Methods

### Cells and viruses

Marc-145 cells, BHK-21 cells and HEK-293T cells were cultured in Dulbecco’s modified Eagle’s medium (DMEM; Gibco, Grand Island, NY, USA) containing 10% fetal bovine serum (FBS, Gibco), 100 U/mL of penicillin and 100 μg/mL streptomycin. Porcine alveolar macrophages (3D4/21) and STING^−/−^ 3D4/21 cells were cultured in RPMI 1640 medium containing 10% FBS with penicillin/streptomycin. All cells were cultured at 37°C in a humidified atmosphere containing 5% CO_2_. The PRRSV strain used in this study was highly pathogenic PRRSV XJ17-5 (PRRSV-2, GenBank ID: MK759853.1) isolated and preserved in our laboratory [37]. The virus was propagated and titrated in Marc-145 cells grown in DMEM supplemented with 2% FBS.

### Antibodies and reagents

The mouse FLAG mAb (HT201), GFP mAb (HT801) and Actin mAb (HC201) were acquired from TransGen Biotech (Beijing, China). The rabbit GFP mAb (AE078), mouse HA mAb (AE008), mouse GAPDH mAb (AC002) and rabbit Histone 3 H3 pAb (A2352) were purchased from ABclonal (Wuhan, Hubei, China). The rabbit STING pAb (19851-1-AP), anti-GORASP2 pAb (10598-1-AP) and Myc pAb (16286-1-AP) were purchased from ProteinTech (Wuhan, Hubei, China). The rabbit mCherry pAb (ab183628) and Calreticulin pAb (ab2907) were acquired from Abcam (Cambridge, Cambridgeshire, UK). The rabbit TBK1 mAb (3504S), phosphorylated-TBK1 (p-TBK1, 5483S) mAb (5483S), IRF3 mAb (11904S), and HA mAb (3724) were acquired from Cell Signaling Technology (Boston, MA, USA). The phosphorylated-IRF3 (p-IRF3, Ser385) rabbit pAb (MA5-14947), Lipofectamine 2000 and Lipofectamine 3000 were purchased from Thermo Fisher Scientific (Sunnyvale, CA, USA). ClonExpress® MultiS One Step Cloning Kit (C113) was purchased from Vazyme Biotech (Nanjing, Jiangsu, China). Protein A/G PLUS-Agarose was purchased from Santa Cruz Biotechnology (sc-2003, CA, USA). The 2’3’-cGAMP was bought from InvivoGen (Hong Kong, China).

### Construction of recombinant expressing plasmids

Total RNA was extracted from HP-PRRSV XJ17-5 using TRIpure reagent, and 22 gene open reading frames (ORFs) of PRRSV were amplified by RT-PCR using the designed primers as shown in Table S1. The PCR products were inserted into the *Nhe*I and *Age*I sites, *Nhe*I and *EcoR*V sites or *Xho*I and *EcoRV* sites of pEGFP-N1 vector. For ten truncated Nsp5 mutants (amino acids 1-101, 1-67, 1-57, 1-47, 1-35, 102-170, 68-170, 58-170, 48-170 and 36-170), the gene fragments were each PCR amplified from pEGFP-N1-Nsp5 using the primers in Table S2 and then cloned into the *Nhe*I and *Age*I sites of pEGFP-N1 vector. For Nsp5 deletion mutants, the two flanking sequences of deletion sites were separately PCR amplified, and then two PCR products were fused by overlapping PCR. The fusion PCR products were cloned into the *Nhe*I and *Age*I sites of pEGFP-N1 vector. All PCR primers used were shown in Table S2. For porcine STING 1-153, the sequence of amino acids 1-153 was amplified by PCR from pmCherry-C1-pSTING plasmid using the primers in Table S3, and cloned into the *Bgl*II and *EcoR*I sites of pmCherry-C1 vector. All the plasmid constructs were confirmed by Sanger DNA sequencing.

pcDNA-DEST-pcGAS-2HA, pcDNA-DEST-pSTING-2HA, pEGFP-C1-pSTING, pmCherry-C1-pSTING, pcDNA-DEST-pSTING-2HA (amino acids 1-190, 191-378, 1-339 and 151-339), pcDNA-Myc-TBK1, pcDNA-Myc-IKKε and p3×FLAG-CMV-7.1-pIRF3 were cloned and stored in our lab.

### Western blotting and Co-immunoprecipitation

The Western blotting was performed as we described previously [21]. Briefly, whole cell protein was extracted and run by SDS-PAGE, and then the protein in gel was transferred to PVDF membrane. The membrane was blocked, incubated with the indicated primary antibodies at 4°C overnight, and then with HRP-labeled anti-rabbit or anti-mouse antibodies (1:1000) (Sangon Biotech, Shanghai, China). The protein signal was visualized by Western blot imaging system using an ECL chemiluminescent detection system (Tanon, Shanghai, China).

For co-immunoprecipitation (Co-IP), the cell lysate from transfected cells in 6-well plate (6-8×10^5^ cells/well) was incubated with 1 μg specific antibodies at 4°C overnight with shaking and further incubated with Protein A/G PLUS-Agarose for 2-3 h. The beads were washed three times with RIPA and eluted with 2×SDS sample buffer. The elution samples together with input lysate controls were both subjected to Western blotting.

### Confocal microscopy

The 3D4/21 cells in 24-well plate were transfected with indicated plasmids and then fixed with 4% paraformaldehyde for 30 min. The fixed cells were permeabilized with 0.5% Triton X-100 for 20 min. Next, the cell nuclei were stained with 4′, 6-diamidino-2-phenylindole (DAPI) for 15 min.

To detect the traffic of STING, mCherry-STING, Nsp5-GFP, Nsp5-GFP (Δ36-47), Nsp5-GFP (Δ58-67), Nsp5-GFP (Δ36-47Δ58-67) or empty vector were transfected into Marc-145 cells and then treated with or without 2’3’-cGAMP. After 24 h, cells were incubated with antibodies (rabbit anti-Calreticulin or rabbit anti-GORASP2) for 1 h at 37℃. Cells were incubated with Alexa Fluor 647-conjugated goat anti-rabbit IgG (H+L) (A32733, Thermo Fisher, USA) for 1 h at 37℃ in the dark and then cell nuclei were stained with DAPI for 15 min. The images were visualized by laser-scanning confocal microscope (LSCM, Leica SP8, Solms, Germany).

### Promoter driven luciferase reporter gene assays

HEK-293T cells grown in 96-well plates (3×10^4^ cells/well) were co-transfected by Lipofectamine 2000 with a firefly luciferase (Fluc) reporter plasmid at 10 ng/well (ISRE-Fluc, ELAM (NF-κB)-Fluc or IFN-β-Fluc) and β-actin *Renilla* luciferase (Rluc) reporter (0.2 ng/well), together with the expression plasmids (20, 50 or 70 ng/well). The total amount of DNA was kept constant by adding vector control plasmid. At 24 h post-transfection, cells were lysed, and samples were assayed as we described previously [38].

### Isolation of cytosolic and nuclear proteins

Cells in 12-well plate were transfected with Nsp5 plasmid for 24 h and then stimulated by transfection with 2’3’-cGAMP for 12 h. After treatment, cytosolic and nuclear proteins were isolated using a Nuclear and Cytoplasmic Protein Extraction Kit (Beyotime, Shanghai, China) following the manufacturer’s instructions and as we described previously [39].

### Construction of recombinant PRRSVs with Nsp5 mutations

To generate PRRSV infectious clones carrying Nsp5 deletions, the full-genome cDNA clones of the pACYC177-CMV-rXJ17-5 and pACYC177-CMV-rXJ17-5-EGFP we constructed previously [25, 40] were used as the skeletons. The fragment 1 (F1) of the two skeleton plasmids were replaced by the F1 with Nsp5 gene deletions. Recombinant PRRSV infectious clones pACYC177-CMV-rXJ17-5/EGFP-Nsp5Δ36-47, pACYC177-CMV-rXJ17-5/EGFP-Nsp5Δ58-67 and pACYC177-CMV-rXJ17-5/EGFP-Nsp5Δ36-47Δ58-67 were obtained, respectively, with the details described in the legend of Fig S2.

### Statistical Analysis

All of the experiments were representative of three similar experiments. The results in bar graphs were presented as the mean ± standard deviation (SD) with three replicates and analyzed using GraphPad Prism 6 software. Statistical analysis was performed using Student’s *t*-test or *ANOVA* where appropriate; *p* < 0.05 was considered statistically significant. In the figures, “*” and “**” denote *p* < 0.05 and *p* < 0.01, respectively.

---

**Fig S1.** The regions of 36-47 and 58-67 amino acids of Nsp5 are the key regions for interaction with STING. 3D4/21 cells were cotransfected with Nsp5-GFP or its mutants and mCherry-STING for 24 h, and then the cells were fixed, permeabilized and examined for cellular co-localization by confocal microscopy. The levels of co-localization between Nsp5 or its truncation mutants and STING were quantified by PCC using ImageJ software. Scale bars are 10 µm.

**Fig S2.** The regions of 36-47 and 58-67 amino acids of Nsp5 are crucial for PRRSV replication. (**A-B**) Schematic diagram of infectious clone recombinant plasmids of rXJ17-5 (A) and rXJ17-5-EGFP (B) carrying the Nsp5 deletions. (**C**) Marc-145 cells were infected with mock, positive controls rPRRSV XJ17-5 and rPRRSV XJ17-5-EGFP, and rPRRSVs carrying Nsp5 deletion mutations, respectively. Cytopathic effects and the GFP signal were observed under fluorescence microscope at 48 hpi.

For the construction of recombinant pACYC177-CMV-rXJ17-5/EGFP-Nsp5Δ36-47, F1-1 was first PCR amplified from the pACYC177-CMV-rXJ17-5-F1 using primers 1 and 5 in Table S4. Second, F1-2 was PCR amplified from pEGFP-N1-Nsp5 using primers 6 and 9 in Table S4. Then, F1-1 and F1-2 PCR products were cloned into the *Pac*I/*Afl*II sites of pACYC177-Stuffer vector using homologous recombination to obtain recombinant pACYC177-CMV-rXJ17-5-F1 (Nsp5Δ36-47). Next, the F1 (Nsp5Δ36-47) was subcloned into the *Pac*I and *Afl*II sites of pACYC177-CMV-rXJ17-5/EGFP vector by T4 ligation to obtain the rXJ17-5/EGFP cDNA clones with 36-47 Nsp5 deletion (pACYC177-CMV-rXJ17-5/EGFP-Nsp5Δ36-47).

For the construction of recombinant pACYC177-CMV-rXJ17-5/EGFP-Nsp5Δ58-67, F1-1 was first PCR amplified from pACYC177-CMV-rXJ17-5-F1 using primers 1 and 7 in Table S4. Second, F1-2 was PCR amplified from pEGFP-N1-Nsp5 using primers 8 and 9 in Table S4. Then, F1-1 and F1-2 PCR products were cloned into the *PacI*/*Afl*II sites of pACYC177-Stuffer vector using homologous recombination to obtain recombinant pACYC177-CMV-rXJ17-5-F1 (Nsp5Δ58-67). Next, the F1 (Nsp5Δ58-67) was subcloned into the *Pac*I and *Afl*II sites of pACYC177-CMV-rXJ17-5/EGFP vector by T4 ligation to obtain the rXJ17-5/EGFP cDNA clones with 58-67 Nsp5 deletion (pACYC177-CMV-rXJ17-5/EGFP-Nsp5Δ58-67).

For the construction of recombinant pACYC177-CMV-rXJ17-5/EGFP-Nsp5Δ36-47Δ58-67, F1-1 was first PCR amplified from pACYC177-CMV-rXJ17-5-F1 using primers 1 and 2 in Table S4. Second, F1-2 was PCR amplified from pEGFP-N1-Nsp5 using primers 3 and 4 in Table S4. Then, F1-1 and F1-2 PCR products were cloned into the *Pac*I/*Afl*II sites of pACYC177-CMV-rXJ17-5/EGFP vector using homologous recombination to directly obtain recombinant pACYC177-CMV-rXJ17-5/EGFP-Nsp5Δ36-47Δ58-67. All the plasmid constructs were confirmed by Sanger DNA sequencing.

The obtained six cDNA clones with Nsp5 deletions and two positive cDNA clones with full length Nsp5 were transfected into BHK-21 cells using Lipofectamine 3000 reagent (ThermoFisher, USA). The cell culture supernatants were obtained at 48 h post transfection and serially passaged on Marc-145 cells to rescue PRRSV deletion mutant virus.

## Data Availability Statement

The authors confirm that the data supporting the findings of this study are available within the article and its supplementary materials.

## Acknowledgement

The work was partly supported by Natural Science Foundation of Shandong Province (ZR2024QC023), National Natural Science Funds (32473040; 32172867), Key Research and Development Project in Shandong Province (Competitive Innovation Platform) (2022CXPT010), Shandong Province Pig Industrial Technology System (SDAIT-08), Taishan Scholars Program of Shandong Province, the 111 Project under Grant D18007, and A Project Funded by the Priority Academic Program Development of Jiangsu Higher Education Institutions (PAPD).

## Author contribution statement

J.Z and J.W conceived and designed the experiments; Y.X, C.C, Q.X, J.Y, Y.Z, P.Z and Wangli Z performed the experiments; S.J, Wanglong Z, N.C and J.W provided the resources; Y.X and J.Z wrote the paper. All authors contributed to the article and approved the submitted version.

